# Biophysical fitness landscape of the SARS-CoV-2 Delta variant receptor binding domain

**DOI:** 10.1101/2022.02.21.481311

**Authors:** Casey Patrick, Vaibhav Upadhyay, Alexandra Lucas, Krishna M.G. Mallela

## Abstract

Among the five known SARS-CoV-2 variants of concern, Delta is the most virulent leading to severe symptoms and increased number of deaths. Our study seeks to examine how the biophysical parameters of the Delta variant correlate to the clinical observations. Receptor binding domain (RBD) is the first point of contact with the human host cells and is the immunodominant form of the spike protein. Delta variant RBD contains two novel mutations L452R and T478K. We examined the effect of single mutations as well as the double mutation on RBD expression in human Expi293 cells, RBD stability using urea and thermal denaturation, and RBD binding to angiotensin converting enzyme 2 (ACE2) receptor and to neutralizing antibodies using isothermal titration calorimetry. Delta variant RBD showed significantly higher expression compared to the wild-type RBD, and the increased expression is due to L452R mutation. Despite their non-conservative nature, none of the mutations significantly affected RBD structure and stability. All mutants showed similar binding affinity to ACE2 and to Class 1 antibodies (CC12.1 and LY-CoV016) as that of the wild-type. Delta double mutant L452R/T478K showed no binding to Class 2 antibodies (P2B-2F6 and LY-CoV555) and a hundred-fold weaker binding to a Class 3 antibody (REGN10987), and the decreased antibody binding is determined by the L452R mutation. These results indicate that the immune escape from neutralizing antibodies, rather than receptor binding, is the main biophysical parameter determining the fitness landscape of the Delta variant RBD and is determined by the L452R mutation.

## Introduction

In late 2019, a novel coronavirus (2019-nCoV), later renamed as severe acute respiratory syndrome coronavirus 2 (SARS-CoV-2), was discovered in Wuhan, China and quickly became the center of the ongoing pandemic coronavirus disease 19 (COVID-19). SARS-CoV-2 enters host cells with its spike protein interacting with the angiotensin converting enzyme 2 (ACE2) located on human host cells.^1–5^ A specific structural region within the spike protein, known as the receptor binding domain (RBD), binds to the ACE2 receptor. SARS-CoV-2 has been shown to continuously mutate in several regions of the spike protein leading to new variants of interest (VOI) and more severe variants of concern (VOC). VOCs in general have been shown to have increased infectivity,^6–9^ enhanced ACE2 binding,^10–12^ escape from the human immune system,^4, 10, 13^, and evade FDA-approved monoclonal antibody therapies.^4, 6, 9^ To date, there have been five known VOCs, which include Alpha, Beta, Gamma, Delta, and Omicron.

Out of all the five VOCs, Omicron is the most transmissible variant, whereas Delta is the most virulent leading to much severe symptoms and increased number of deaths. Delta variant arose from the B.1.617 lineage, and is specifically labeled as variant B.1.617.2. It was first identified in India in March 2021,^14^ and has since been accounted for the majority of COVID-19 deaths worldwide.^15^ Delta variant has been shown to have higher viral titers in COVID-19 patients compared to previous variants.^15–17^ Prior to the emergence of Omicron variant, increased breakthrough infections of COVID-19 in vaccinated patients have been attributed to the Delta variant.^16–19^ A single dose of vaccination was found to be only 33% effective in protecting against the Delta variant as opposed to 48.7% against the Alpha variant, and two vaccination doses were only 88% effective for the Delta variant compared to 93.7% for the Alpha variant.^20^

Delta variant introduces several mutations in the N-terminal domain (NTD), RBD, and the furin cleavage site of the spike protein that makes it distinct from the unmutated, wild-type (WT) virus.^21^ Unlike previous variants, which have had mutations that have been predicted by *in vitro* evolution through biophysical parameters such as ACE2 binding, the mutations in the Delta variant, particularly in the RBD, have not previously been predicted to lead to a more dangerous variant.^12, 22^ Delta RBD contains two mutations which change the characteristic nature of the amino acid: a hydrophobic amino acid leucine mutated to a positively charged amino acid arginine at position 452 (L452R) and an uncharged amino acid threonine mutated to a positively charged lysine at position 478 (T478K) in the primary structure of the protein.^6, 9, 14, 23, 24^ None of these two mutations are part of the RBDs of previously discovered VOCs that include Alpha (N501Y), Beta (K417N/E484K/N501Y), or Gamma (K417T/E484K/N501Y). The newly discovered Omicron VOC RBD contains only the T478K mutation and not the L452R mutation.^25^

Analyzing the biophysical parameters that determine the fitness landscape of viruses is of considerable interest in recent years, particularly in the case of HIV, influenza, dengue, hepatitis C, and retroviruses.^26–31^ In the case of SARS-CoV-2 RBD, we along with others have recently shown that increased receptor binding, escape from neutralizing antibodies, and maintaining protein structure, stability, and expression despite the non-conservative nature of amino acid mutations are important parameters that direct the natural selection of mutations and determine the biophysical fitness landscape of emerging variants.^10, 32–34^ These biophysical analyses were done on Alpha, Beta, and Gamma VOCs before the Delta variant has emerged. Whether the two novel mutations of the Delta RBD, which were not part of the previous VOCs, follow similar natural selection principles is not clear. This study examined the effect of the two single amino acid mutations L452R, T478K and the double mutant L452R/T478K on the biophysical properties (structure, stability, receptor binding, and binding to neutralizing antibodies) of the RBD. Our results indicate that the Delta RBD does not show increased ACE2 receptor binding unlike the previous VOCs (Alpha, Beta, and Gamma), but shows increased expression, consistent with increased spike protein expression and increased viral titers in Delta patients, and escapes multiple neutralizing antibodies. Increased expression and antibody escape is solely determined by the L452R mutation.

## Results

### L452R mutation enhances Delta variant RBD expression

VOCs including Delta have been found to have decreased levels of neutralization titers in both vaccinated and unvaccinated individuals.^15–17, 35^ In addition, Delta variant COVID-19 patients have viral titers ten times higher than that of the other variants.^15^ A plausible explanation is that the mutations in the Delta variant might have a selective advantage in terms of increased expression of viral proteins over the wild-type virus. Higher quantities of the viral proteins could allow for more virus particles to be created.^36^ In order to test if this clinical observation could be correlated with the increased expression of RBD, which is a major part of the spike protein, expression of the wild-type (WT) RBD and its Delta mutants (L452R, T478K, and L452R/T478K) was tested in modified HEK293 cells (Expi293). After 48 hours of post-transfection, secreted proteins in the supernatants were analyzed using SDS-PAGE (Figure 1A) and the expression levels were quantified as a ratio of the mutant over WT RBD (Figure 1B). Both the L452R single mutant and the Delta double mutant L452R/T478K showed ∼70% higher expression compared to WT RBD (Figure 1B). No significant differences in expression were observed for the T478K mutant compared to the WT RBD. These results indicate that the L452R mutation is responsible for the increased expression of Delta variant RBD and possibly the spike protein expression.

**Figure 1.**
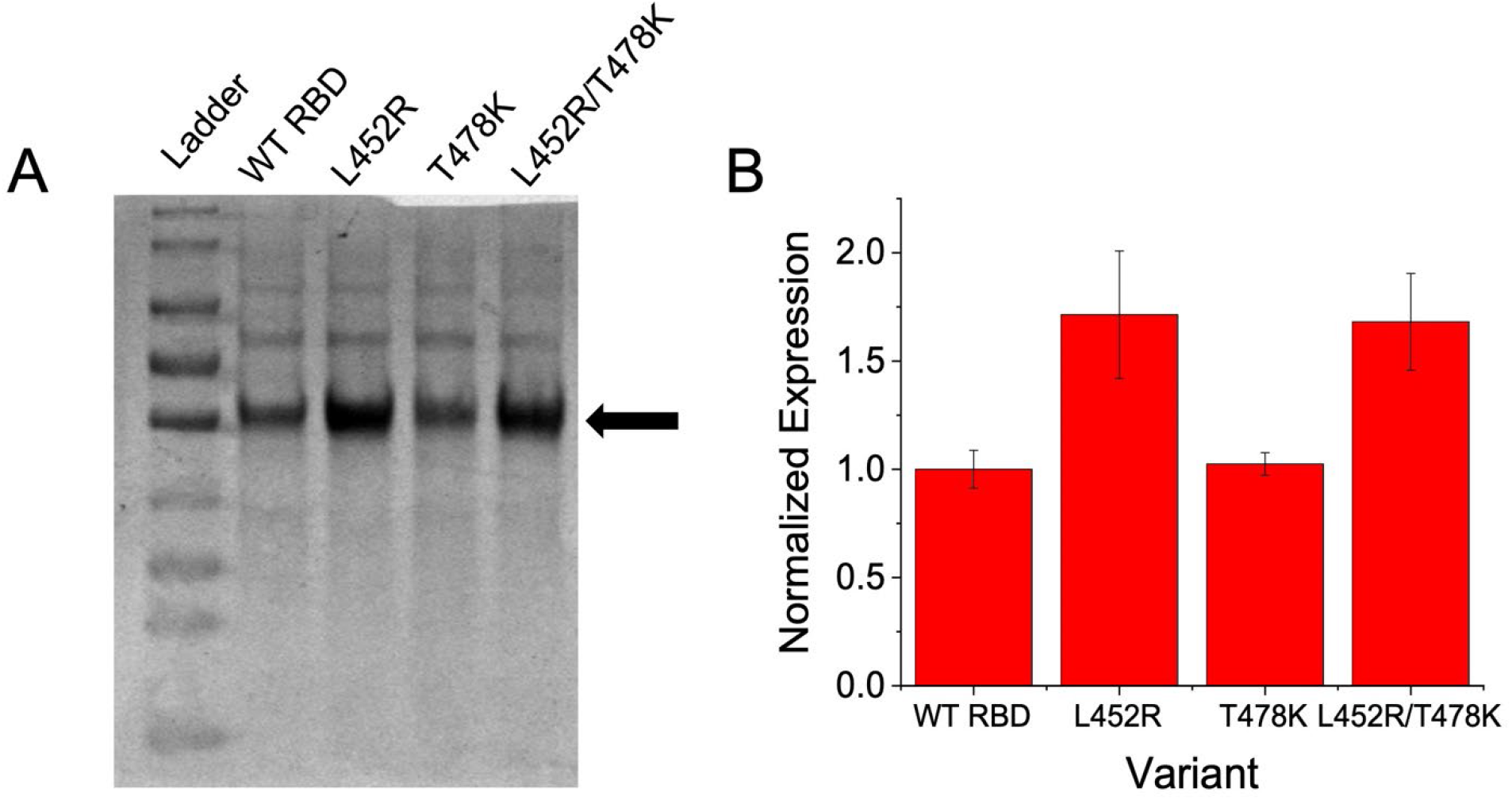
Expression differences between WT RBD and its Delta mutants. (A) SDS-PAGE showing relative expression of RBD variants 48 hours after transfection. Ladder represents the molecular markers (from top to bottom: 180, 130, 100, 70, 55, 40, 35, 25, and 15 kDa, respectively). (B) Normalized relative expression of RBD variants. Results in panel A were repeated in triplicate to obtain the data shown in panel B.

### None of the Delta mutations significantly affect global protein structure of RBD

Both mutations L452R and T478K are non-conservative mutations where one type of amino acid is mutated to another type of amino acid with differing physical properties. Such mutations tend to destabilize proteins if the amino acid prior to mutation is involved in stabilizing the protein structure. To test the effect of mutations on RBD structure, we used far-UV circular dichroism and fluorescence spectroscopy. Figure 2A shows SDS-PAGE of purified proteins, and the single bands on the gel show the high purity of protein samples used for biophysical analyses reported in this manuscript. Figure 2B shows the far-UV circular dichroism (CD) and Figure 2C shows the intrinsic protein fluorescence spectra of the WT RBD and its Delta single mutants and the double mutant. Spectra of the WT match those reported in the literature.^10, 37–39^ More importantly, none of the mutations caused significant changes in the spectra, implying that the Delta mutations do not affect the global protein structure of RBD.

**Figure 2.**
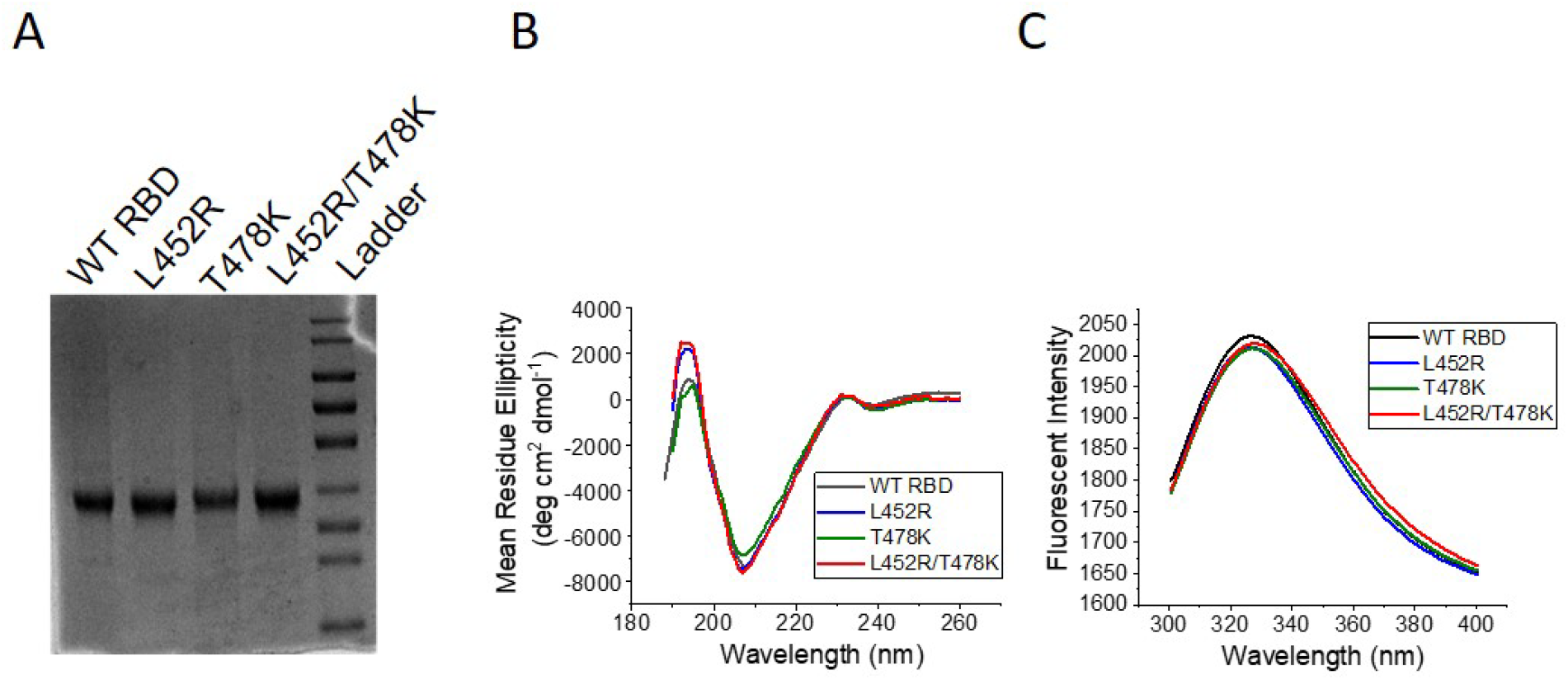
(A) SDS-PAGE of purified RBD constructs. Ladder represents the molecular markers (from top to bottom: 180, 130, 100, 70, 55, 40, 35, 25, and 15 kDa, respectively). (B) Far-UV circular dichroism spectra of WT RBD and its Delta mutants. (C) Intrinsic fluorescence spectra of WT RBD and its Delta mutants.

### None of the Delta mutations significantly affect RBD stability

Protein stability could provide valuable insight into both the viability and flexibility of proteins, and has been shown to play a big role in the fitness of viruses.^27, 40^ To evaluate how the Delta mutations alter the RBD stability, both thermal and urea denaturation melts were utilized. Change in protein structure with increase in temperature (Figure 3) was fit to a two-state unfolding model (Equation 1 in Materials and Methods) to obtain the midpoint melting temperature (T_m_) of the proteins. Since the thermal melts are not reversible, T_m_ values can only be used as a qualitative measure of protein stability.^10, 41^ Table 1 lists the mean fit parameters obtained from three independent batches of protein expression. Compared to WT RBD, which showed a T_m_ of 56.1 ± 0.7°C, L452R displayed a similar stability of 56.5 ± 0.2°C, while T478K displayed a slightly decreased T_m_ of 54.1 ± 0.1°C. However, the Delta double mutant L452R/T478K exhibited a T_m_ of 56.6 ± 0.2°C, similar to that of the WT (Figure 3 & Table 1).

**Figure 3.**
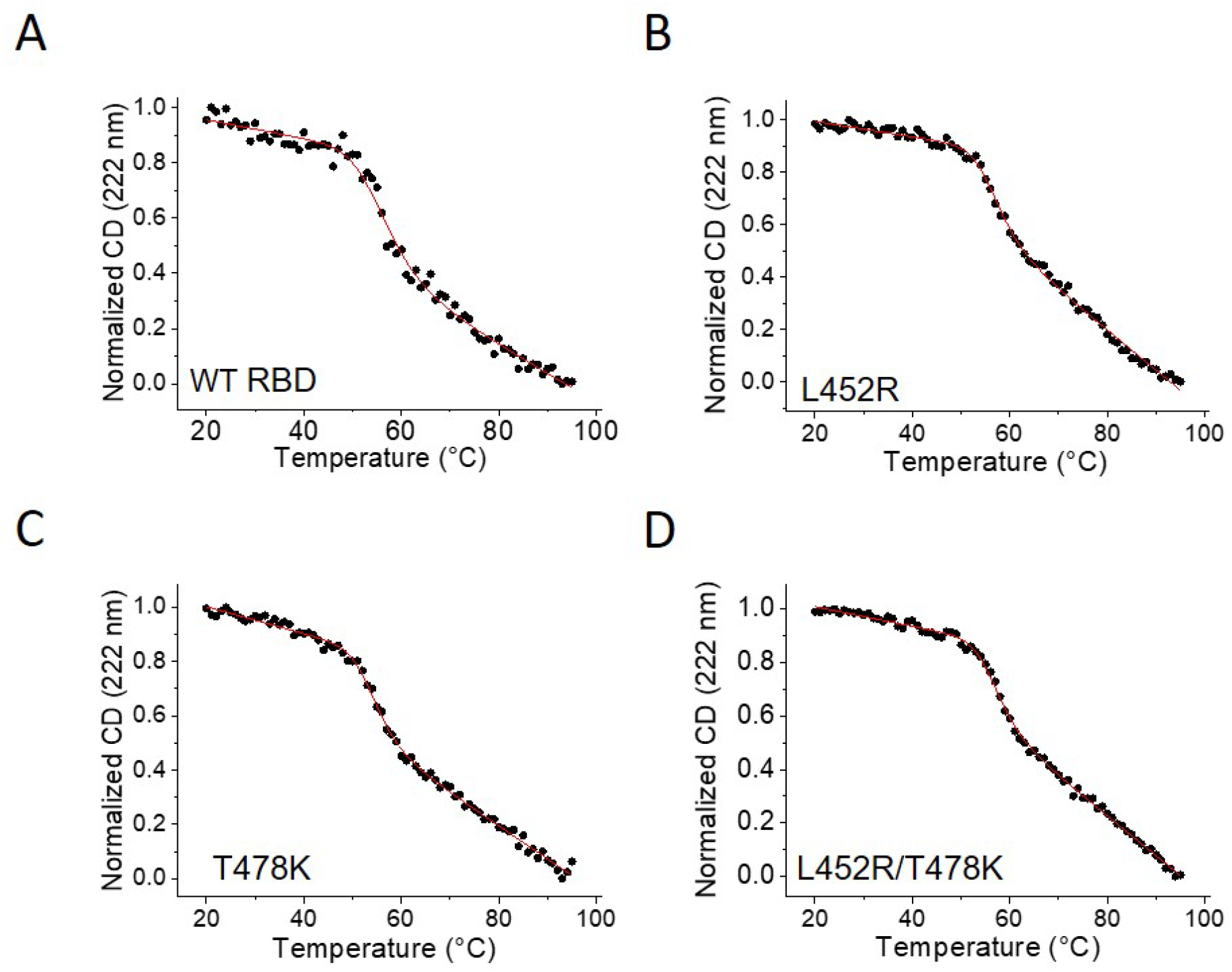
Normalized thermal denaturation of WT RBD and its Delta mutants monitored using far-UV circular dichroism.

**Table 1.** Thermal denaturation fit values for WT RBD and its Delta mutants.

| **RBD Variant** | **T<sub>m</sub> (°C)** | **ΔH (kcal/mol)** |
| --- | --- | --- |
| WT | 56.1 ± 0.7 | -1.7 ± 0.3 |
| L452R | 56.5 ± 0.2 | -2.4 ± 0.2 |
| T478K | 54.1 ± 0.1 | -2.3 ± 0.2 |
| L452R/T478K | 56.6 ± 0.2 | -2.6 ± 0.2 |

Urea denaturation melts of RBD variants are completely reversible (Figure 4). The native signal showed a large change with denaturant concentration, which might indicate partial unfolding and non-2-state unfolding behavior that needs to be further probed. However, since we observed only a single sigmoidal transition, denaturant melts were fit to a 2-state unfolding model (Equation 2 in Materials and Methods) to obtain Gibbs free energy of unfolding in the absence of denaturant (ΔG°_unf_) and the slope of linear variation of ΔG_unf_ with urea concentration (m-value) for each variant. Table 2 lists the mean fit parameters obtained from three independent batches of protein expression. WT RBD showed a ΔG°_unf_ of 8.1 ± 0.3 kcal/mol with a m-value of -1.24 ± 0.06 kcal/mol/M [urea]. Both single mutants L452R and T478K showed similar stability as that of the WT RBD. L452R displayed a ΔG°_unf_ of 8.1 ± 0.2 kcal/mol with a m-value of -1.43 ± 0.04 kcal/mol/M [urea], while T478K showed ΔG°_unf_ and m-values of 7.9 ± 0.5 kcal/mol and -1.37 ± 0.09 kcal/mol/M [urea], respectively. Delta double mutant L452R/T478K was also found to have similar stability as that of the WT RBD, with a ΔG°_unf_ of 8.6 ± 0.3 kcal/mol and an m-value of -1.53 ± 0.06 kcal/mol/M [urea] (Table 2). These equilibrium stability values obtained from urea denaturant melts (Table 2) agree quite well with the trends observed with thermal denaturation melts (Table 1), and indicate that none of the Delta mutants significantly affect RBD stability.

**Figure 4.**
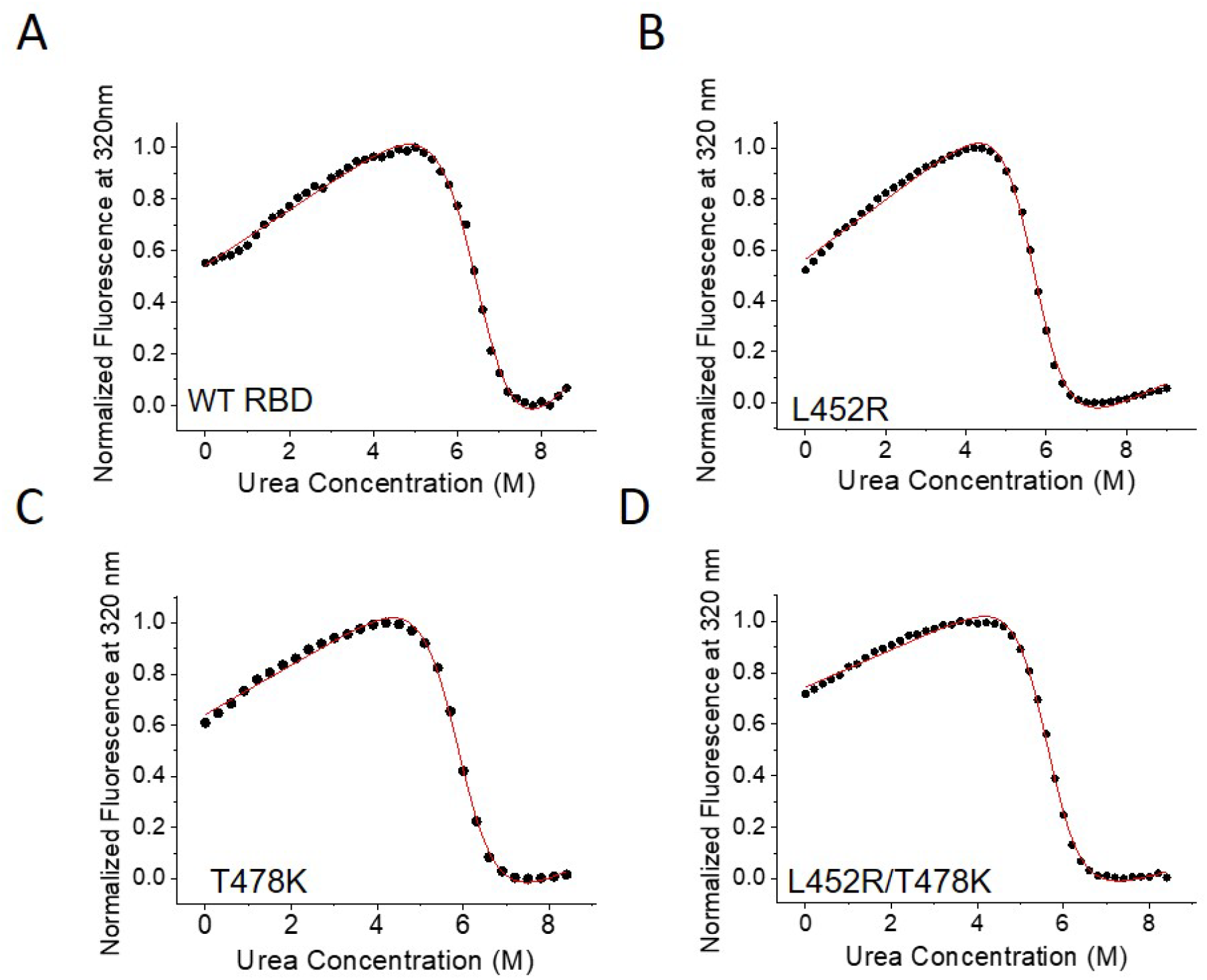
Normalized urea denaturation melts of WT RBD and its Delta mutants monitored using intrinsic protein fluorescence.

**Table 2.** Chemical Denaturation fit values for WT RBD and its Delta mutants.

| **RBD Variant** | **ΔG<sub>unf</sub><sup>o</sup> (kcal/mol)** | **m-value (kcal/mol/M [Urea])** |
| --- | --- | --- |
| WT | 8.1 ± 0.3 | -1.24 ± 0.06 |
| L452R | 8.1 ± 0.2 | -1.43 ± 0.04 |
| T478K | 7.9 ± 0.5 | -1.37 ± 0.09 |
| L452R/T478K | 8.6 ± 0.3 | -1.53 ± 0.06 |

### Delta mutations do not show increased affinity for ACE2 receptor

Since SARS-CoV-2 enters host cells with its RBD binding to ACE2, the relative binding affinity of the RBD can play a major role in how variants are evolving. An increase in the affinity of the Delta variant to ACE2 could allude to a potential mechanism where the VOC allows more viral entry into host cells. Previous VOCs (Alpha, Beta, and Gamma) have shown enhanced ACE2 binding compared to the WT RBD.^10^ Location of the two amino acid mutations L452R and T478K in the Delta variant RBD with respect to ACE2 binding interface is shown in Figure 5A. None of the two mutations are part of the ACE2 interface. Figure 6 shows isothermal titration calorimetry (ITC) thermograms for the WT RBD and Delta mutants, and Table 3 lists the average fit parameters from three independent batches of protein expression. WT RBD binding to ACE2 shows a typical exothermic reaction with a K_d_ of 10.0 ± 3.1 nM and a ΔH value of -11.8 ± 0.2 kcal/mol. None of the Delta mutations significantly altered the binding affinity of RBD for ACE2. Both L452R and T478K mutants and the Delta double mutant L452R/T478K displayed similar K_d_ values of 6.2 ± 3.7 nM, 17.1 ± 4.8 nM, and 10.8 ± 3.3 nM, respectively, similar to that of the WT RBD (Table 3).

**Figure 5.**
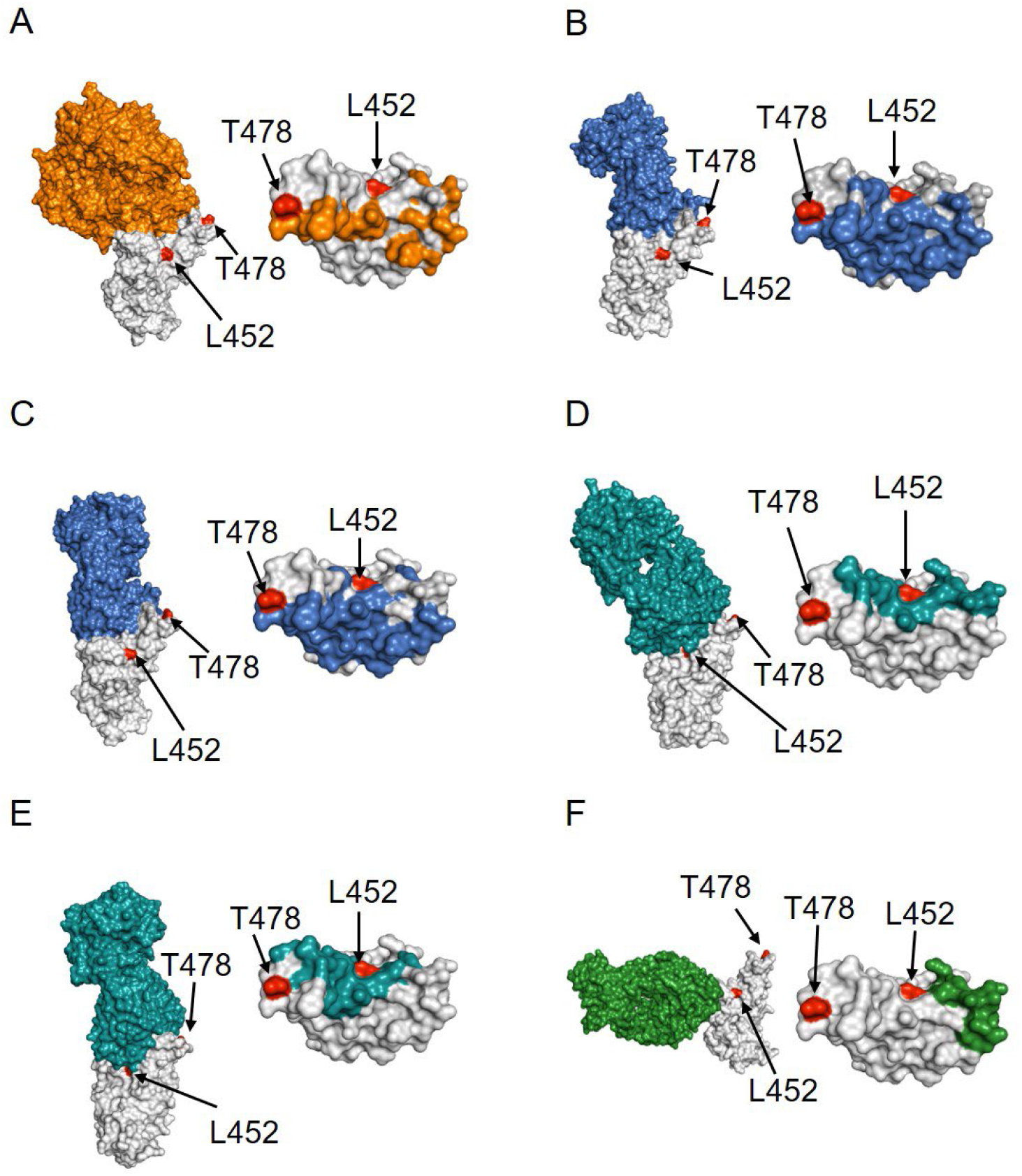
Structural analysis of the location of Delta mutants L452 and T478 (red colored) with respect to RBD (gray colored) complexes with (A) ACE2 receptor (orange colored; PDB ID 6moj), (B) Class 1 antibody CC12.1 (blue colored; PDB ID 6xc2), (C) FDA-approved Class I therapeutic antibody LY-CoV016 (blue colored; PDB ID 7c01), (D) Class 2 antibody P2B-2F6 (teal colored; PDB ID: 7bwj), (E) FDA-approved Class 2 antibody LY-CoV555 (teal colored; PDB ID: 7kmg) and (F) FDA-approved Class 3 antibody REGN10987 (green colored; PDB ID 6xdg). Left panels show the complex structures and right panels show the location of the interacting residues in RBD.

**Figure 6.**
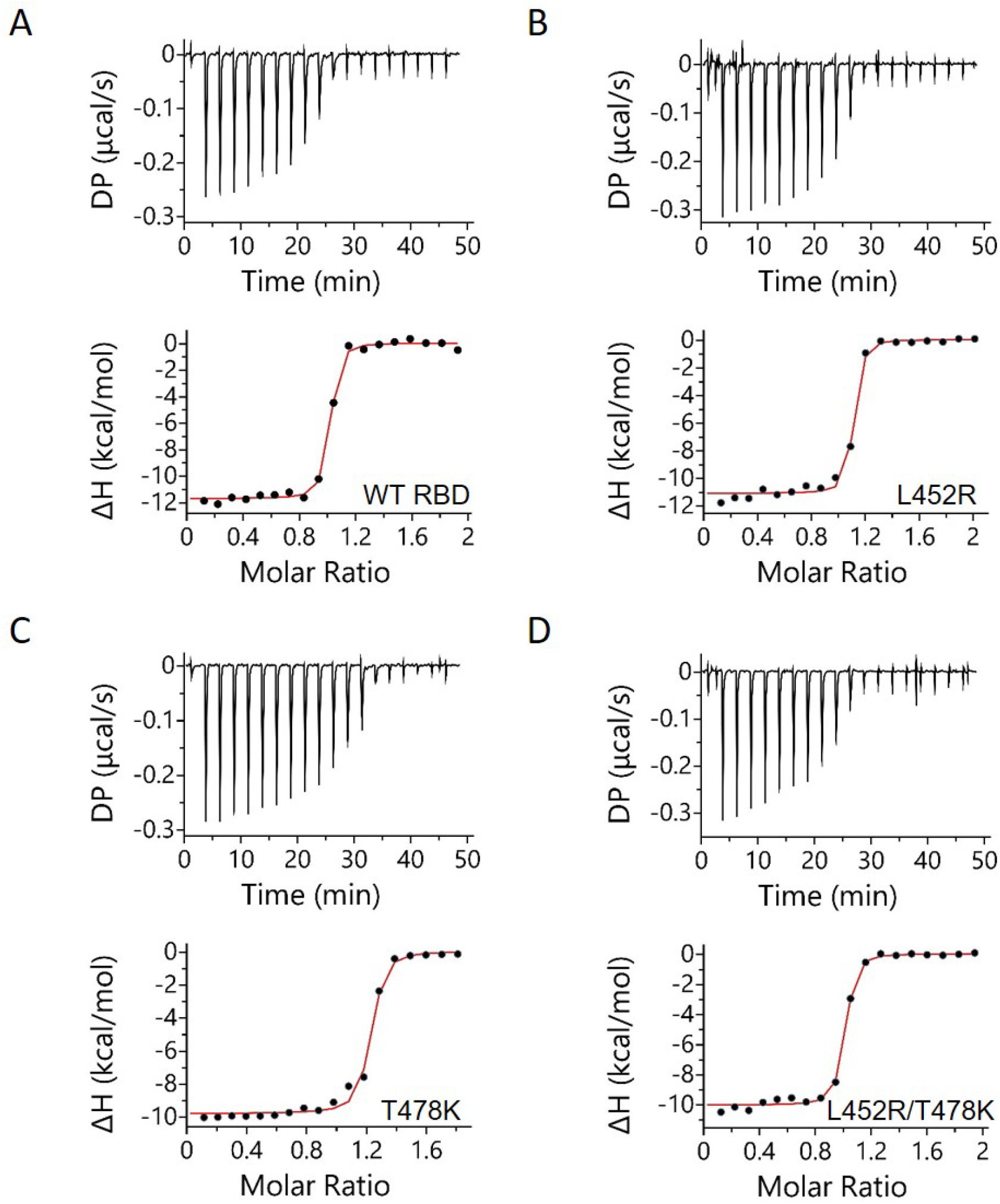
ITC analysis of WT RBD and its Delta mutants binding to ACE2. Top panels represent the raw differential power vs. time thermographs, while bottom panels represent the integrated heat plots.

**Table 3.** Thermodynamic parameters of WT RBD and its Delta mutants binding to ACE2 receptor.

| **RBD Variant** | **K<sub>d</sub>**<br>**(nM)** | **N** | **ΔH (kcal/mol)** | **ΔG (kcal/mol)** | **-TΔS**<br>**(kcal/mol)** |
| --- | --- | --- | --- | --- | --- |
| WT | 10.0 ± 3.1 | 1.0 ± 0.0 | -11.8 ± 0.2 | -10.7 ± 0.2 | 1.0 ± 0.3 |
| L452R | 6.2 ± 3.7 | 1.0 ± 0.0 | -11.3 ± 0.2 | -11.1 ± 0.3 | 0.3 ± 0.4 |
| T478K | 17.1 ± 4.8 | 1.0 ± 0.1 | -10.0 ± 0.2 | -10.4 ± 0.2 | - 0.4 ± 0.3 |
| L452R/T478K | 10.8 ± 3.3 | 0.9 ± 0.0 | -10.1 ± 0.2 | -10.7 ± 0.2 | - 0.6 ± 0.3 |

### Delta RBD mutations do not escape Class 1 antibodies

Clinical observations associated with Delta variant could be related to SARS-CoV-2 escaping the human immune system. Neutralizing antibodies against SARS-CoV-2 RBD have been found to belong to four major classes depending on the mechanism of action and the location of their epitopes on the RBD.^42, 43^ SARS-CoV-2 spike protein is a trimer in its native state and exists in multiple conformations, mainly RBD in “up” position which is accessible for binding to ACE2 or in “down” position in which RBD is buried and not accessible for ACE2 binding.^44–46^ Class 1 antibodies bind to RBD in the up conformation and compete with ACE2 binding. Class 2 antibodies bind to RBD both in the up or down conformations, and their epitope partially overlaps with the ACE2 binding site and hence compete against ACE2 binding. Class 3 antibodies bind to RBD in both up or down positions with their epitope on RBD far away from ACE2 binding site, and hence can neutralize the RBD through an uncompetitive mechanism. Class 4 antibodies are relatively rare as their epitope is close to the hinge region connecting RBD to the rest of the spike protein, which is relatively buried compared to other epitopes, and none of the VOCs contain mutations in this region. Since the first step in neutralization is binding of antibodies to RBD, we examined how the Delta mutations affect RBD binding to the three major classes of antibodies.

One of the first Class 1 antibodies that was identified from patients recovered from WT SARS-CoV-2 infection was CC12.1.^47^ Location of the two Delta mutants L452 and T478 in RBD with respect to the CC12.1 binding interface is shown in Figure 5B. Binding of WT RBD and its Delta mutants to CC12.1 single chain variable fragment (ScFv) was measured using ITC (Figure 7), and the average thermodynamic parameters obtained from fitting the data from three independent batches of protein expression was included in Table 4. All proteins showed similar exothermic binding profiles. WT RBD binds to CC12.1 with a K_d_ of 23.9 ± 5.7 nM. Both Delta single mutants L452R and T478K as well as the double mutant L452R/T478K either bind with a similar affinity (similar K_d_ value for T478K) or with a higher affinity (lower K_d_ values for L452R and L452R/T478K), implying that the Delta variant RBD does not escape from Class 1 antibodies. ITC binding data is also consistent with the location of these two residues L452 and T478 with respect to the epitope of CC12.1 on RBD (Figure 5B). The two mutations in the Delta variant RBD are located far from the CC12.1 epitope, and do not affect the interactions between RBD and CC12.1.^48, 49^

**Figure 7.**
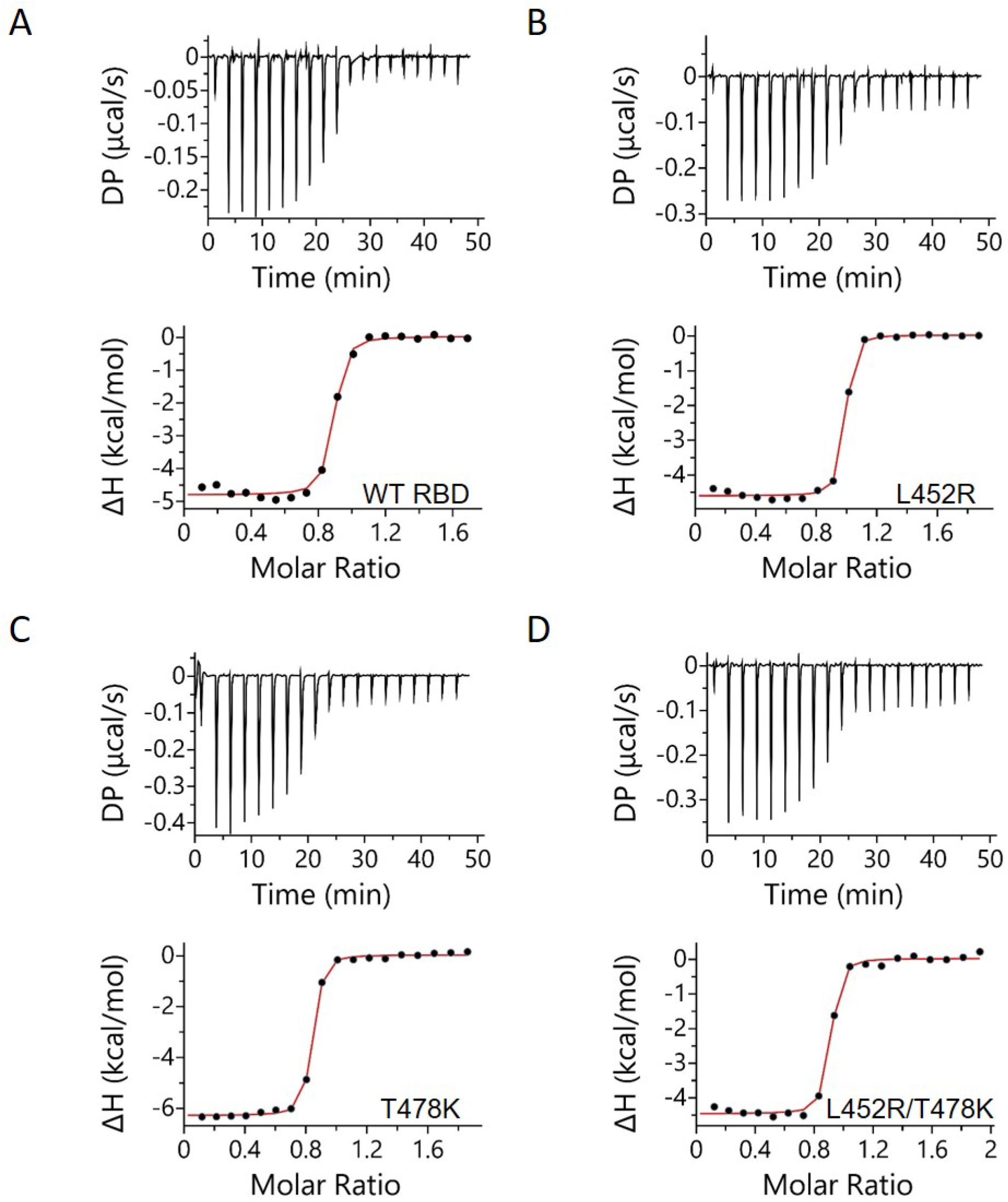
ITC analysis of WT RBD and its Delta mutants binding to Class 1 antibody CC12.1 ScFv. Top panels represent the raw differential power vs. time thermographs, while bottom panels represent the integrated heat plots.

**Table 4.**
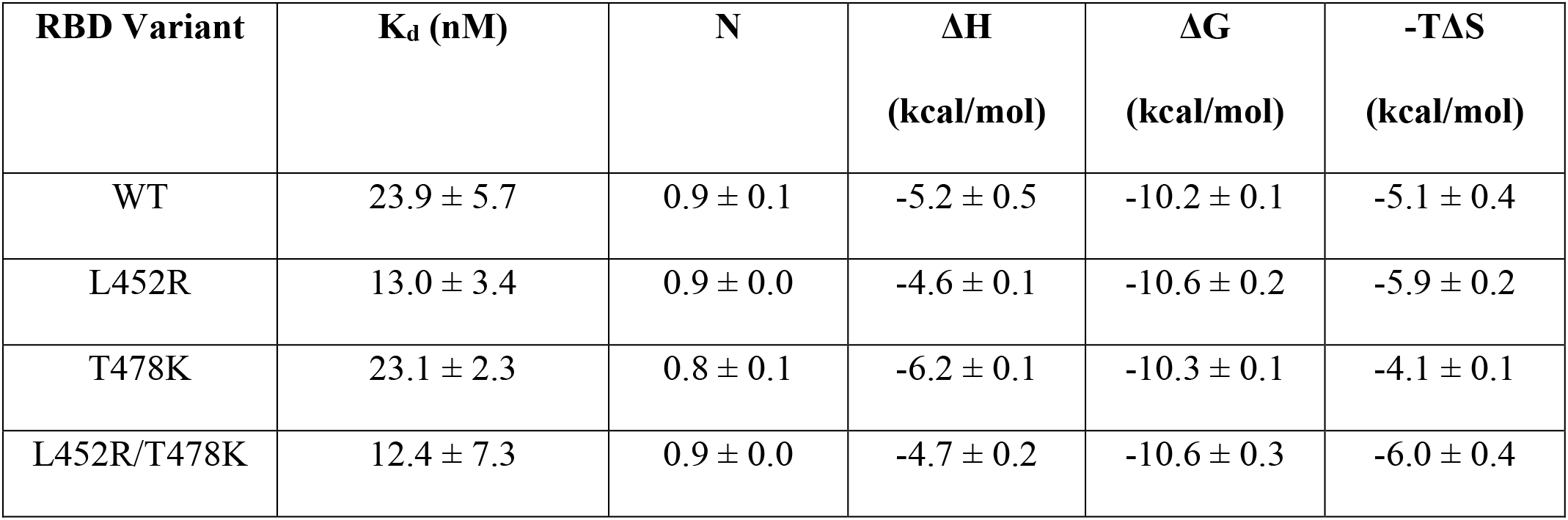
Thermodynamic parameters of WT RBD and its Delta mutants binding to Class 1 antibody CC12.1 ScFv.

| **RBD Variant** | **K<sub>d</sub> (nM)** | **N** | **ΔH**<br>**(kcal/mol)** | **ΔG**<br>**(kcal/mol)** | **-TΔS**<br>**(kcal/mol)** |
| --- | --- | --- | --- | --- | --- |
| WT | 23.9 ± 5.7 | 0.9 ± 0.1 | -5.2 ± 0.5 | -10.2 ± 0.1 | -5.1 ± 0.4 |
| L452R | 13.0 ± 3.4 | 0.9 ± 0.0 | -4.6 ± 0.1 | -10.6 ± 0.2 | -5.9 ± 0.2 |
| T478K | 23.1 ± 2.3 | 0.8 ± 0.1 | -6.2 ± 0.1 | -10.3 ± 0.1 | -4.1 ± 0.1 |
| L452R/T478K | 12.4 ± 7.3 | 0.9 ± 0.0 | -4.7 ± 0.2 | -10.6 ± 0.3 | -6.0 ± 0.4 |

As of January 2022, FDA approved four therapeutic antibodies for emergency use authorization (EUA): Eli Lilly’s LY-CoV016 (Etesevimab) and LY-CoV555 (Bamlanivimab), and Regeneron’s REGN10933 (Casirivimab) and REGN10987 (Imdevimab). LY-CoV016 and REGN10933 are Class 1 antibodies whose epitopes are very similar on RBD.^50, 51^ LY-CoV555 is a Class 2 antibody, whereas REGN10987 is a Class 3 antibody. To determine whether Delta variant escapes FDA-approved Class 1 antibodies, we tested WT RBD and its Delta mutants binding to LY-CoV016 in ScFv format. Figure 5C shows the location of the two residues L452 and T478 with respect to the RBD interface with LY-CoV016. Figure 8 shows the ITC binding data for the single and double mutants. All interactions displayed exothermic binding profiles. Table 5 lists the average parameters obtained from fitting the ITC data from three independent batches of protein expression to a one-site binding model. WT RBD binds to LY-CoV016 with a K_d_ of 49.3 ± 10.1 nM. Both single mutants L452R and T478K and the double mutant L452R/T478K bind to LY-CoV016 with a stronger affinity, implying that none of the Delta mutations escape LY-CoV016. This is consistent with the location of the two residues L452 and T478 with respect to the RBD binding interface with LY-CoV016 (Figure 5C), and also consistent with binding of Delta mutants to another Class 1 antibody CC12.1 described above (Figure 7 and Table 4).

**Figure 8.**
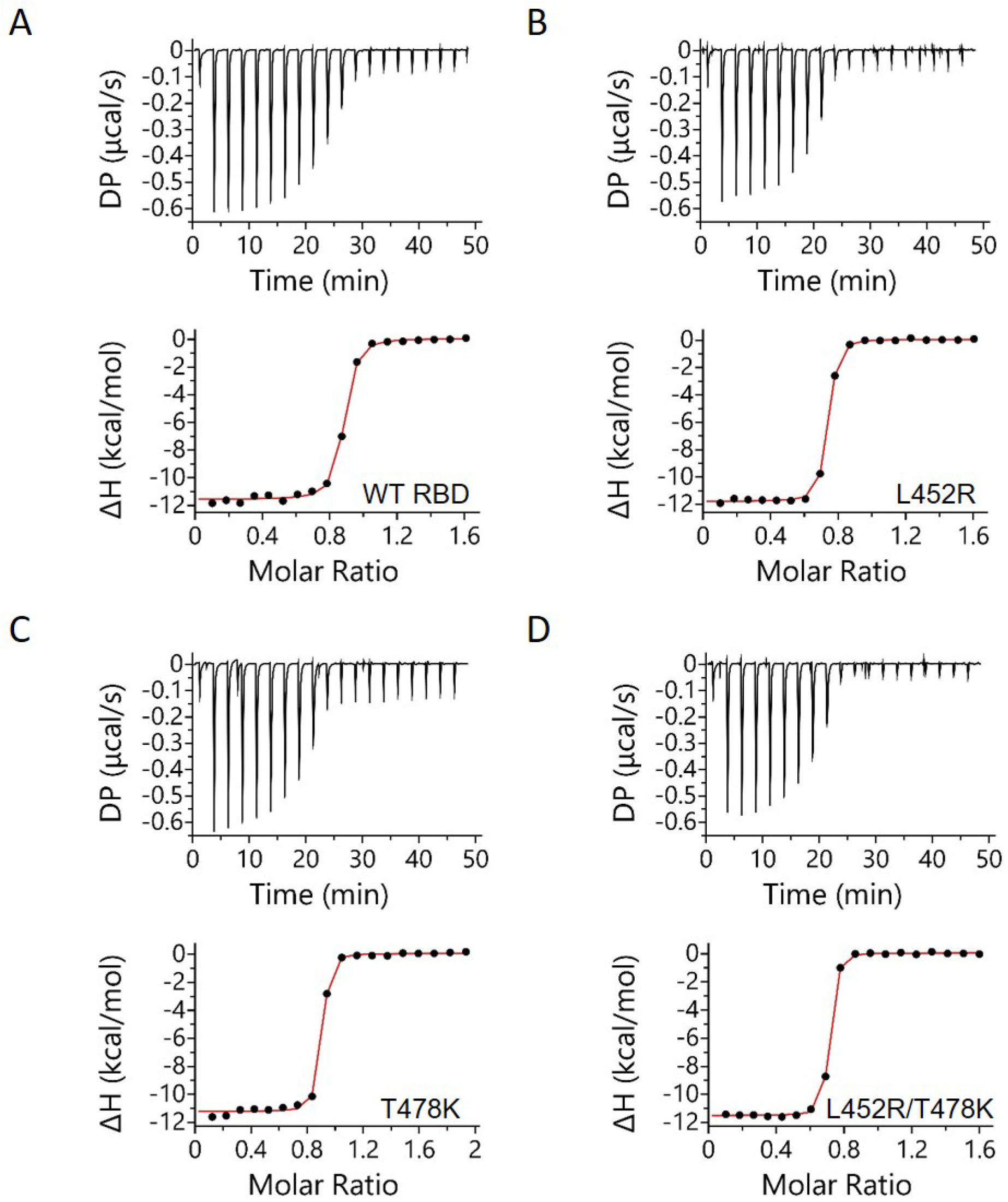
ITC analysis of WT RBD and its Delta mutants binding to FDA-approved Class 1 therapeutic antibody LY-CoV016 ScFv. Top panels represent the raw differential power vs. time thermographs, while bottom panels represent the integrated heat plots.

**Table 5.** Thermodynamic parameters of WT RBD and its Delta mutants binding to FDA-approved Class 1 therapeutic antibody LY-CoV016 ScFv.

| **RBD Variant** | **K<sub>d</sub> (nM)** | **N** | **ΔH**<br>**(kcal/mol)** | **ΔG**<br>**(kcal/mol)** | **-TΔS**<br>**(kcal/mol)** |
| --- | --- | --- | --- | --- | --- |
| WT | 49.3 ± 10.1 | 0.9 ± 0.0 | -11.7 ± 0.1 | -9.8 ± 0.1 | 1.9 ± 0.2 |
| L452R | 23.4 ± 2.8 | 0.8 ± 0.1 | -11.9 ± 0.1 | -10.2 ± 0.1 | 1.7 ± 0.1 |
| T478K | 15.4 ± 3.0 | 0.8 ± 0.1 | -11.4 ± 0.1 | -10.5 ± 0.1 | 0.9 ± 0.2 |
| L452R/T478K | 10.8 ± 1.4 | 0.8 ± 0.1 | -12.0 ± 0.4 | -10.7 ± 0.1 | 1.3 ± 0.4 |

### L452R mutation determines Delta variant escape from Class 2 antibodies

While the Delta variant does not evade Class 1 antibodies, a large amount of clinical data suggests that neutralizing antibodies discovered against WT SARS-CoV-2 are not effective in neutralizing emerging VOCs.^52^ Similar to CC12.1, P2B-2F6 is one of the first neutralizing antibodies discovered in recovered COVID-19 patients.^53, 54^ P2B-2F6 is a Class 2 antibody.^42, 43^ We examined whether Delta variant escapes from Class 2 antibodies by determining the binding of WT RBD and its Delta mutants to P2B-2F6 ScFv. Location of the two residues L452 and T478 in RBD with respect to its binding interface with P2B-2F6 is shown in Figure 5D. Figure 9 shows the ITC binding curves and Table 6 lists the thermodynamic parameters obtained from fitting ITC data from three independent batches of protein expression to a one-site binding model. Both WT RBD and T478K show similar K_d_ values of 81.4 ± 6.2 nM and 80.5 ± 6.3 nM, respectively. However, L452R resulted in a complete loss of binding (Figure 9). Similar results were observed for the Delta double mutant L452R/T478K. These results indicate that the Delta variant escapes Class 2 antibodies, and is determined by the L452R mutation.

**Figure 9.**
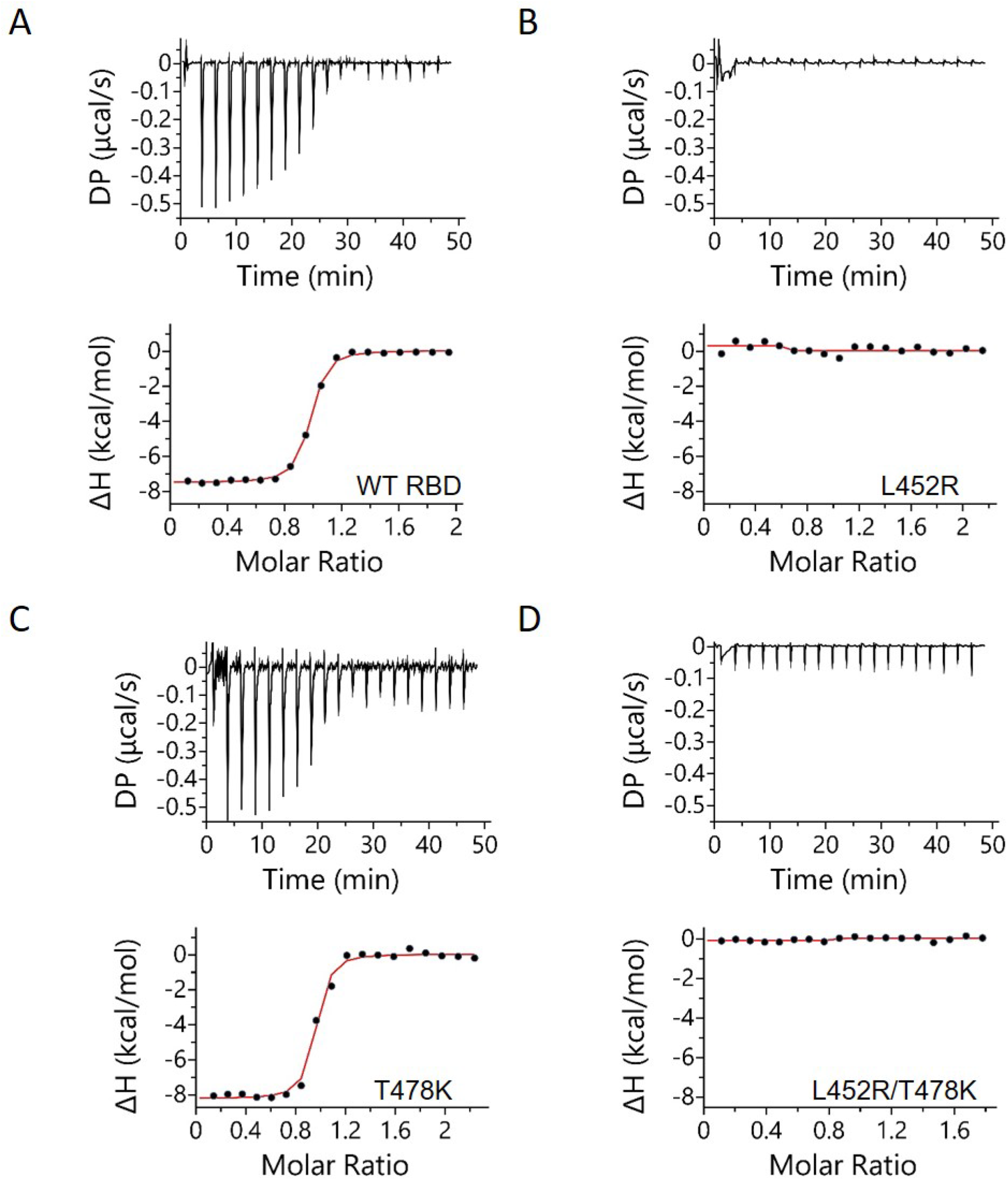
ITC analysis of WT RBD and its Delta mutants binding Class 2 antibody P2B-2F6 ScFv. Top panels represent the raw differential power vs. time thermographs, while bottom panels represent the integrated heat plots.

**Table 6.** Thermodynamic parameters of WT RBD and its Delta mutants binding to Class 2 antibody P2B-2F6 ScFv.

| **RBD Variant** | **K<sub>d</sub>**<br>**(nM)** | **N** | **ΔH**<br>**(kcal/mol)** | **ΔG**<br>**(kcal/mol)** | **-TΔS**<br>**(kcal/mol)** |
| --- | --- | --- | --- | --- | --- |
| WT | 81.4 ± 6.2 | 0.9 ± 0.0 | -8.0 ± 0.5 | -9.5 ± 0.1 | -1.5 ± 0.5 |
| L452R | No binding | - | - | - | - |
| T478K | 80.5 ± 6.3 | 0.9 ± 0.0 | -8.7 ± 0.4 | -9.5 ± 0.0 | -0.8 ± 0.4 |
| L452R/T478K | No binding | - | - | - | - |

We further examined whether Delta variant escapes FDA-approved Class 2 antibody LY-CoV555.^55, 56^ Location of the two residues L452 and T478 in RBD with respect to its interface with LY-CoV555 is shown in Figure 5E. Figure 10 shows the ITC binding curves for WT RBD and its Delta mutants, and Table 7 lists the mean thermodynamic parameters obtained from fitting ITC data from three independent batches of protein expression to one-site binding model. WT RBD binds to LY-CoV555 ScFv with a K_d_ of 3.8 ± 1.9 nM. T478K mutant shows a similar binding affinity to LY-CoV555 with a K_d_ of 9.0 ± 4.6 nM. This was however not the case for the L452R mutation and the Delta double mutant L452R/T478K. Both showed no binding to LY-CoV555 (Figure 10). These results indicate that Delta variant escapes Class 2 antibodies and is determined by the L452R mutation, which is consistent with escape from another Class 2 antibody P2B-2F6 described above (Figure 8 and Table 6).

**Figure 10.**
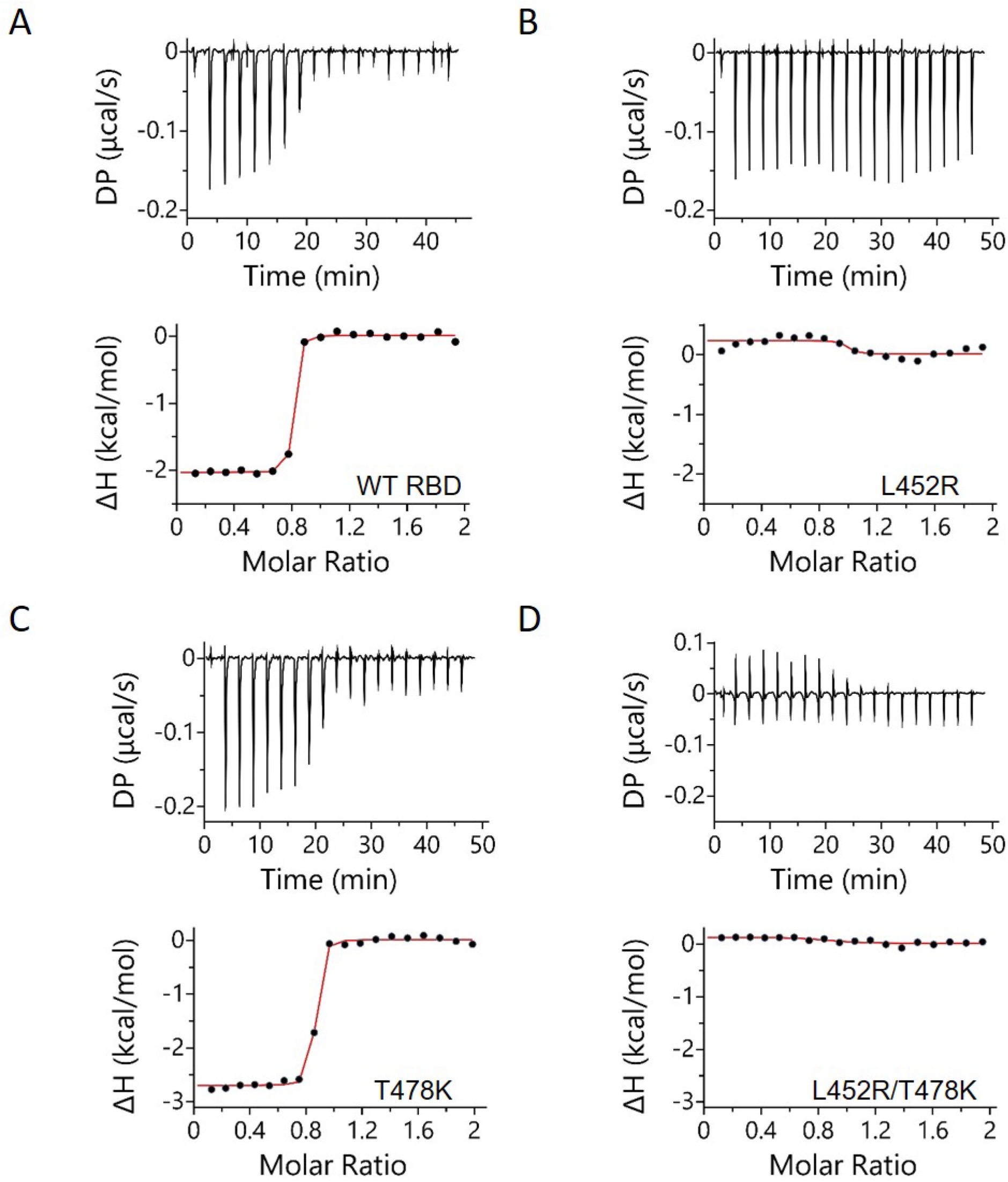
ITC analysis of WT RBD and its Delta mutants binding to FDA-approved Class 2 therapeutic antibody LY-CoV555 ScFv. Top panels represent the raw differential power vs. time thermographs, while bottom panels represent the integrated heat plots.

**Table 7.** Thermodynamic parameters of WT RBD and its Delta mutants binding to FDA-approved Class 2 therapeutic antibody LY-CoV555 ScFv.

| **RBD Variant** | **K<sub>d</sub>**<br>**(nM)** | **N** | **ΔH**<br>**(kcal/mol)** | **ΔG**<br>**(kcal/mol)** | **-TΔS**<br>**(kcal/mol)** |
| --- | --- | --- | --- | --- | --- |
| WT | 3.8 ± 1.9 | 0.8 ± 0.1 | -2.0 ± 0.1 | -11.3 ± 0.3 | -9.3 ± 0.3 |
| L452R | No binding | - | - | - | - |
| T478K | 9.0 ± 4.6 | 0.8 ± 0.0 | -2.7 ± 0.1 | -10.8 ± 0.3 | -8.1 ± 0.3 |
| L452R/T478K | No binding | - | - | - | - |

### L452R mutation determines Delta variant escape from Class 3 antibodies

We also examined whether Delta variant escapes Class 3 antibodies. FDA-approved antibody therapeutics contain a Class 3 antibody REGN10987. Location of the two residues L452 and T478 in RBD with respect to its interface with REGN10987 is shown in Figure 5F. Figure 11 shows the ITC binding curves of WT RBD and its Delta mutants, and Table 8 lists the mean thermodynamic parameters obtained from fitting ITC data from three independent batches of protein expression to a one-site binding model. Both WT RBD and T478K mutant showed similar binding affinity with K_d_ values of 34.3 ± 8.1 nM and 15.9 ± 1.9 nM, respectively. This was not the case for both the L452R mutant and the Delta double mutant L452R/T478K. Both proteins showed a ∼100 fold weaker binding affinity with K_d_ values of 2,700 ± 1,400 nM for the L452R mutant and 1,500 ± 400 nM for the Delta double mutant (Table 8). These results indicate that the Delta variant escapes Class 3 antibodies, and the escape is determined by the L452R mutation.

**Figure 11.**
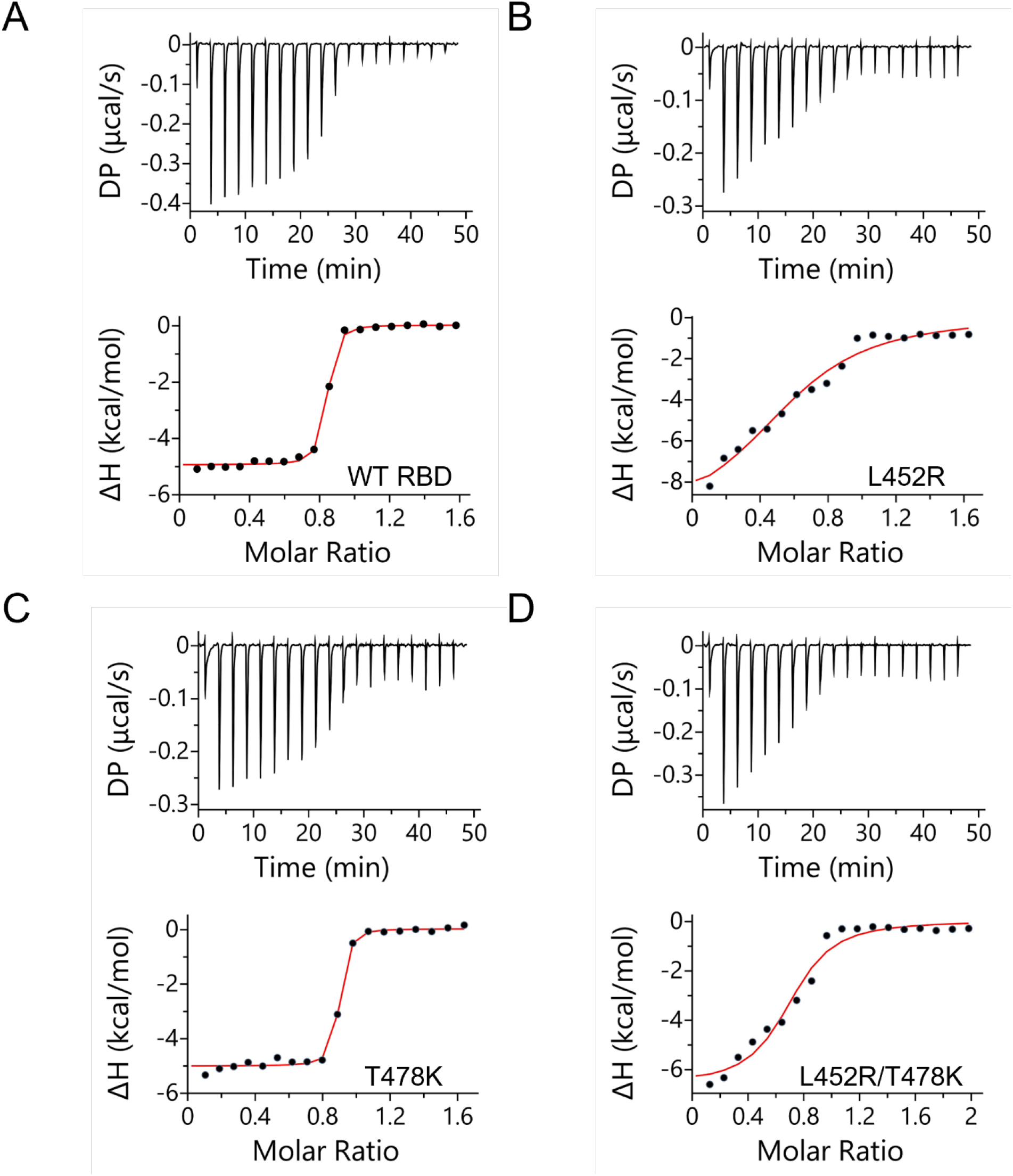
ITC analysis of WT RBD variant and its Delta mutants binding to FDA-approved Class 3 therapeutic antibody REGN987 ScFv. Top panels represent the raw differential power vs. time thermographs, while bottom panels represent the integrated heat plots.

**Table 8.**
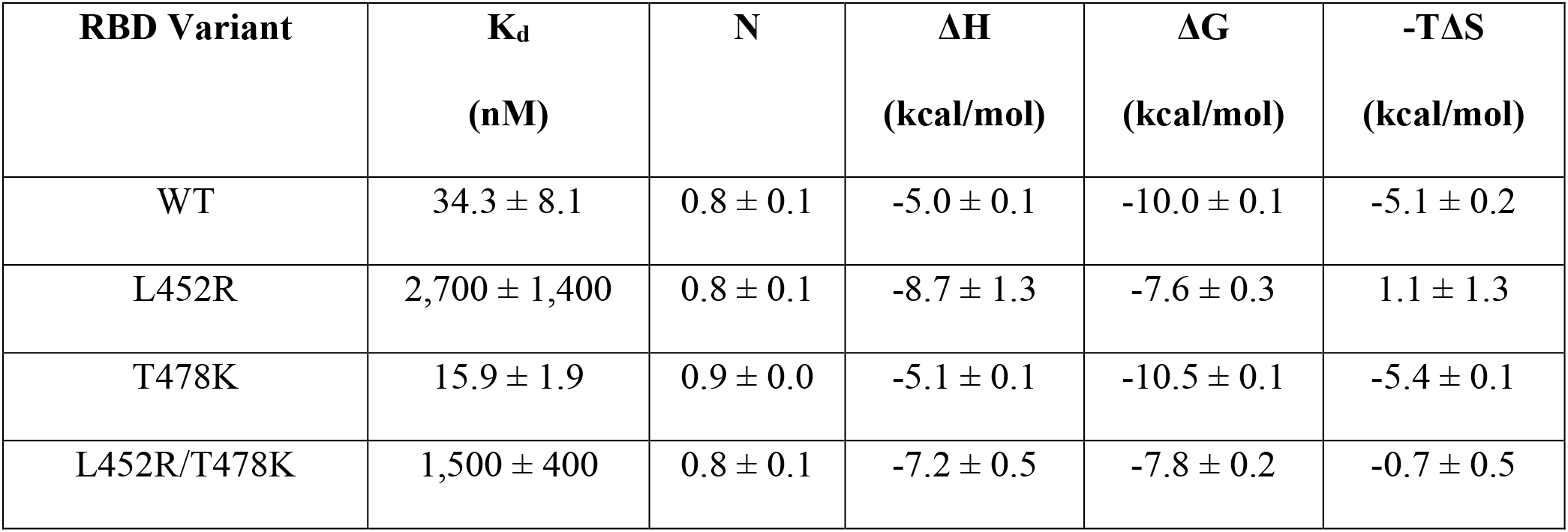
Thermodynamic parameters of WT RBD and its Delta mutants binding to FDA-approved Class 3 therapeutic antibody REGN10987 ScFv.

## Discussion

SARS-CoV-2 Delta variant has shown to adapt unlike any other previous VOCs (Alpha, Beta, and Gamma). It is the VOC which is responsible for more severe symptoms and maximum number of deaths compared to other VOCs including Omicron. Previous studies were unable to predict the two novel mutations in the Delta variant with high probability before the emergence of the variant. None of the two mutations L452R and T478K are part of Alpha, Beta, and Gamma VOCs, and only T478K (and not L452R) is present in the recently discovered Omicron VOC. In this manuscript, we examined the effect of these two novel mutations on the biophysical fitness landscape of Delta variant RBD.

Delta VOC displays unique biophysical characteristics unlike the previous VOCs Alpha, Beta, and Gamma. Table 9 lists the summary of biophysical parameters we have examined. Delta mutations do not significantly alter the binding affinity of RBD towards the ACE2 receptor (Figure 6 & Table 3). While a common belief is that VOCs should result in increased binding to ACE2, which would correlate with increased viral entry, data on Delta variant shows that this cannot be a ubiquitous thought. VOC having no effect on ACE2 binding affinity is unique to the Delta variant, as all previously VOCs showed increased affinity to ACE2.^10, 34^ This is primarily because of the presence of N501Y mutation in previous VOCs that is responsible for increased ACE2 binding.^57, 58^ Delta VOC does not contain N501Y mutation, whereas all other VOCs including Omicron contains the N501Y mutation. In addition, neither L452 nor T478 have a direct interaction with ACE2 (Figure 5A). ACE2 binding has been one of the most common factors when attempting to predict emerging VOCs,^12^ which would explain why the two novel RBD mutations that resulted in the Delta variant have not been predicted by earlier studies. Thus, it is important to consider a more robust system for predicting variants that is not so heavily weighted towards ACE2 binding, as our results show that immune escape rather than receptor binding determines the fitness of Delta VOC. Since the full-length spike protein exists in multiple conformations with RBDs in up or down positions,^13, 59^ measured binding affinity for isolated RBD towards ACE2 represents the upper value of the binding affinity. Any conformation with RBD in down position in equilibrium will only decrease the relative population of RBDs in up conformation, and hence will result in decreased affinity of the complete spike protein towards ACE2.

**Table 9.**
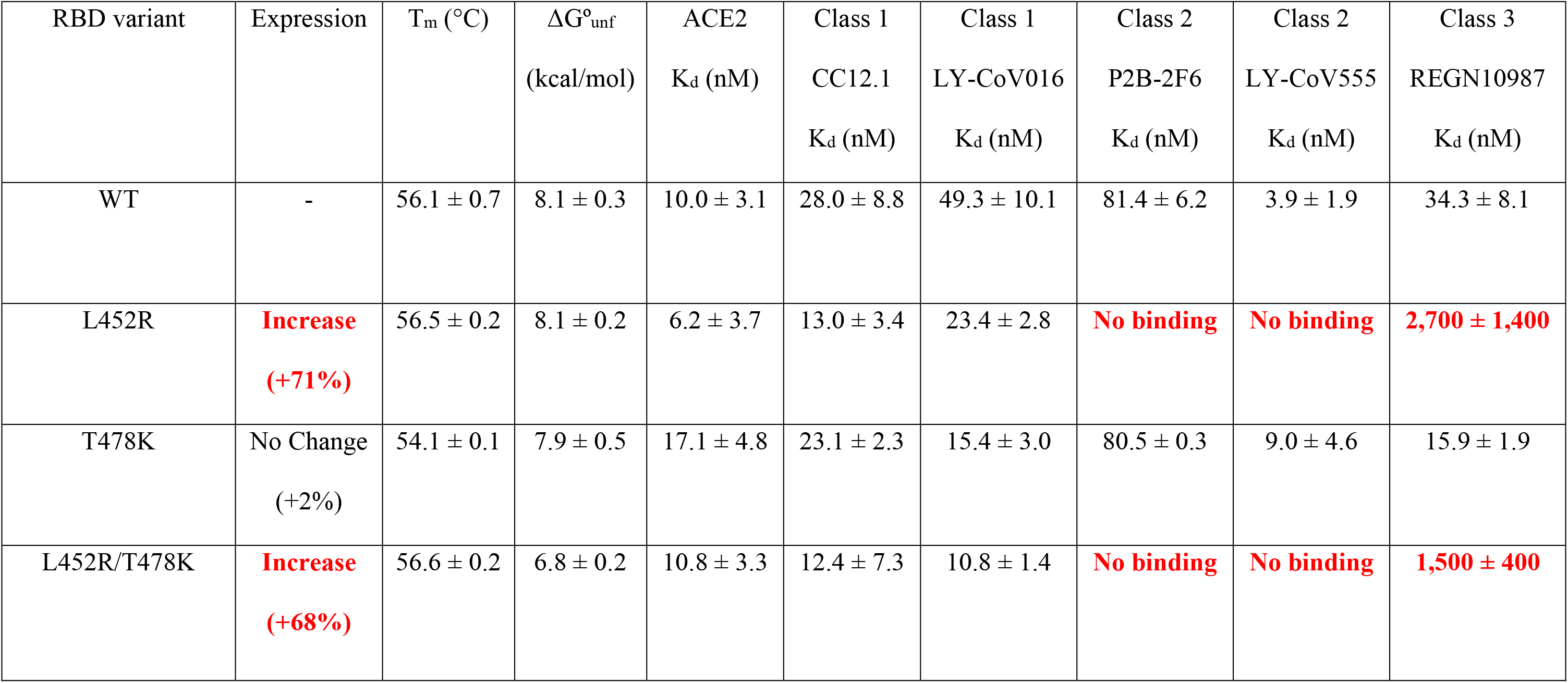
Biophysical parameters determining the fitness of SARS-CoV-2 Delta variant RBD.

Compared to other VOCs, patients contracted with Delta have increased viral titers.^15, 60^ Results show that the two Delta mutations do not affect either the secondary (far-UV CD; Figure 2B) or tertiary structure (intrinsic protein fluorescence of aromatic sidechains; Figure 2C) of the RBD (Figure 5). Additionally, the two mutations L452R and T478K did not alter the stability of the RBD when measured by both thermal denaturation (Figure 3 & Table 1) or urea denaturation (Figure 4 & Table 2) experiments. However, Delta RBD showed ∼70% higher expression in human Expi293 (modified HEK293) cells compared to the WT RBD (Figure 1). Similar high expression of RBD was not seen in the case of Alpha, Beta, and Gamma VOCs.^10^ Increased expression of RBD is entirely determined by the L452R mutation (Figure 1 and Table 9), and this mutation is not present in other VOCs. One single mutation increasing the protein expression by 70% is very rare in protein literature in general. Increased expression of viral proteins can lead to increased viral titers, although whether L452R results in increased expression of the complete spike protein needs to be examined. If that is the case, relative expression levels of viral protein mutants need to be considered as a criterion in predicting the emergence of future VOCs.

Another important factor to consider when evaluating viral fitness is the ability of mutations to escape neutralizing antibodies generated by human immune system in response to WT SARS-CoV-2 infection, vaccination, or those authorized by FDA for treating infected patients. Determining whether the variants escape FDA-approved antibodies, which are mainly derived either from patients recovered from WT SARS-CoV-2 infection (Eli Lilly) or from humanized mice models (Regeneron), will tell us whether the current therapies work against the emerging variants or new antibody therapies have to be developed. Neutralizing antibodies have been broadly classified into different classes depending on the location of their epitopes on RBD.^42, 43^ Locations of mutations in the structures of RBD-antibody complexes (Figure 5) and analyzing the stabilizing interactions in which they participate can sometimes predict which antibodies the VOCs can escape. However, experiments have to confirm such predictions based on protein structure because long-range mutation effects and the dynamics of various protein regions can also play a critical role in protein function. The two mutations L452R and T478K are far away from the binding interface of two Class 1 antibodies we examined (CC12.1 (Figure 5B) & FDA-approved Eli Lilly Class 1 antibody LY-CoV016 (Figure 5C)), and both mutations do not participate in any stabilizing interactions of RBD with Class 1 antibodies. Consistently, Delta mutants did not escape Class 1 antibodies (Figure 7, Figure 8, Table 4, Table 5 & Table 9). In contrast, L452 stabilizes the RBD interactions with Class 2 antibodies by forming a hydrophobic cluster with I103 and V105 of the variable heavy chain of P2B-2F6 and I54 and L55 residues of the variable heavy chain of LY-CoV555. Replacing the hydrophobic residue leucine with a positively charged arginine in the middle of these hydrophobic clusters is expected to destabilize the RBD interactions with Class 2 antibodies. Accordingly, after L452R mutation, RBD does not bind to Class 2 antibodies (Figure 9, Figure 10, Table 6, Table 7 & Table 9). In the case of RBD binding to Class 3 antibodies (Figure 5F), L452 does not form any direct contacts with the antibody REGN10987, but neighbors the epitope composed of residue 450 in RBD.^61, 62^ Change of a hydrophobic amino acid leucine with a positively charged residue arginine next to the epitope is expected to change the electrostatic nature of RBD interface with REGN10987, and thus might explain the decrease in binding affinity by ∼100-fold upon L452R mutation (Figure 11 & Table 9). T478 residue is far away from any of the binding interfaces and do not participate in any stabilizing inter-molecular interactions, and hence do not contribute to the immune escape potential of the Delta variant (Table 9).

Above results indicate that the Delta variant has evolved towards escape from Class 2 and Class 3 antibodies, rather than enhancing the receptor binding or escape from Class 1 antibodies. Class 1 antibodies bind to RBD only in up conformation where RBD is accessible to ACE2 binding, whereas Class 2 and Class 3 antibodies bind to RBD irrespective of whether it is in up conformation (accessible to ACE2) or down conformation (inaccessible to ACE2).^43^ Escape from Class 2 and Class 3 antibodies mainly contributes to escape from polyclonal plasma,^50^ which might be more important for virus survival than escape from Class 1 antibodies that target only a sub-population of the spike protein trimers with their RBDs in up conformation. Further, since Class 2 and Class 3 antibodies can bind to RBD irrespective of whether it is in up (ACE2 accessible) or down (ACE2 inaccessible) conformation, these antibodies can recognize adjacent RBDs in the spike trimer and once bound they can lock RBDs in down conformation thereby restricting binding to ACE2;^63^ hence, the virus escaping from Class 2 and Class 3 antibodies might be more relevant for the spike protein of the variants to bind to ACE2 leading to increased infection. In terms of the efficacy of the current FDA-approved antibody therapies, both Eli Lilly’s Class 1 antibody LY-CoV016 and Regeneron’s Class 1 antibody REGN10933 should be effective in neutralizing the Delta variant, since the Delta mutations did not affect RBD binding to Class 1 antibodies (Table 9). However, Eli Lilly’s Class 2 antibody LY-CoV555 will be completely ineffective in neutralizing the Delta variant as L452R completely abolished RBD binding to a Class 2 antibody, whereas Regeneron’s Class 3 antibody REGN10987 will be much less effective and require much higher concentration as the RBD binding affinity is reduced by ∼100 fold upon Delta mutations (Table 9).

The immune escape and high expression capabilities of the SARS-CoV-2 Delta variant requires a necessity for robust therapeutic options. As the virus adapts, every VOC has shown increased immune escape potential. Thus, it is necessary for the vaccination rates to continue to rise in order to combat the emergence of future VOCs. Simultaneously, it is necessary for improved monoclonal antibody therapeutics to be developed to combat future variants. Delta VOC is clearly distinct from other VOCs. Our previous work has shown that other VOCs can escape Class 1 antibodies, while expression is not significantly different from that of the unmutated WT.^10^ Delta does not escape Class 1 antibodies, and shows higher protein expression. In addition, all previously studied variants showed enhanced ACE2 binding which is not the case for the Delta variant. These results also point to the fact that the virus is still under continuous evolution, as none of the VOCs is still able to escape all classes of neutralizing antibodies. Any combination of mutations that confer immune escape potential to SARS-CoV-2 against all classes of neutralizing antibodies will be of a major concern. In terms of applicability of our results to the new Omicron variant, Omicron still shows higher ACE2 binding,^64^ probably because of the presence of N501Y mutation. Since Omicron lacks L452R mutation, its escape from Class 2 and Class 3 neutralizing antibodies^65^ might be conferred by a different set of amino acid mutations. If most transmissible Omicron picks up the most virulent Delta mutations resulting in a Deltacron variant, it could turn out to be a more dangerous variant.

## Materials and Methods

### Cloning

Sequences for RBD and human ACE2 were obtained from Uniprot (RBD ID: P0DTC2 hACE2 ID: Q9BYF1). Sequences for neutralizing antibodies were obtained from Research Collaboratory for Structural Bioinformatics (RCSB) Protein Data Bank (PDB). CC12.1, LY-CoV016, P2B-2F6, LY-CoV555, and REGN 10987 had PDB IDs of 6XC2, 7C01, 7BWJ, 7KMG, and 6XDG respectively. Antibody constructs were designed in ScFv format, linking the heavy chain (V_H_) and light chain (V_L_) via a glycine serine linker. Final ScFv constructs were V_H_ – (GGGGS)_3_–V_L._ All sequences were codon optimized for expression in mammalian cells by Twist Biosciences. Final constructs included the protein of interest, SUMOstar protein attached to a His-tag, and a human immunoglobulin heavy chain secretory sequence from 5’ to 3’ position. RBD variants were created by site-directed mutagenesis. Sequences were cloned into a pcDNA3.4-TOPO vector. Expi293 HEK cells were transfected at a concentration of 3 x 10^6^ cells/mL with polyethyleneimine. Proteins were expressed over a 5-day period, and the culture media was centrifuged and the supernatant was filter sterilized with a 0.22 µm PVDF filter. The expression levels for individual mutants were compared after running the culture supernatants on SDS-PAGE, staining with Coomassie blue R-250 dye, and quantifying the band intensities corresponding to the target protein using the ImageLab software from Bio-Rad.

### Purification

All proteins were purified using a Ni-NTA column, and the protein was eluted with 200 mM imidazole. The eluted protein was dialyzed to remove imidazole and stored in a buffer consisting of 50 mM Tris, 200 mM NaCl, pH 8.0. The proteins were cleaved using SUMOstar protease at 4°C overnight. Proteins were once again passed through a Ni-NTA column, and the digested proteins were collected in the flow-through and wash. Proteins were dialyzed into a buffer solution of 50 mM sodium phosphate, 20 mM NaCl, pH 7.0. Purity was confirmed by SDS-PAGE.

### CD Spectroscopy

An Applied Photophysics Chirascan Plus spectrometer was used to record the CD spectra and chemical denaturation melts for all variants. CD Spectra were obtained for each RBD variant from 190 nm to 260 nm. Protein spectra were recorded with a 0.5 mm cuvette at 5 µM protein concentration in a buffer consisting of 10 mM sodium phosphate, 4 mM NaCl, pH 7.0. Spectra were recorded every 1 nm wavelength and averaged over 2 seconds. Runs were repeated 5 times and averaged.

Thermal denaturation melts were performed using an Applied Photophysics Chirascan Plus spectrometer. All experiments were performed in a 0.5 mm cuvette at a protein concentration of 20 µM in buffer containing 50 mM sodium phosphate, 20 mM NaCl, pH 7.0. Spectra were recorded 222 nm and averaged over 2 seconds, and the temperature scan was recorded at a rate of 1°C. Data was plotted and analyzed using the equation^41^

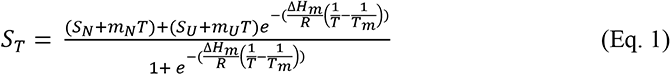

where S_T_ is the measured signal as a function of temperature T, S_N_ and S_U_ are the signals corresponding to the native and unfolded baselines, m_N_ and m_U_ are the slopes of linear dependence of S_N_ and S_U_ with temperature, T_m_ is the midpoint melting temperature, ΔH_m_ is the enthalpy change at the T_m_, R is the universal gas constant, and T is the absolute temperature in Kelvin, respectively.

A CCD detector was used with the Applied Photophysics Chirascan Plus spectrometer to record the chemical denaturation spectra. An excitation wavelength of 280 nm was used, and fluorescence emission was recorded in a 1 cm cuvette at 2 µM protein concentration in a buffer containing 50 mM sodium phosphate, 20 mM NaCl, pH 7.0. Urea was utilized as the denaturant for equilibrium protein unfolding measurements, and was dissolved in 50 mM sodium phosphate, 20 mM NaCl, pH 7.0 buffer. The concentration of the urea solution was determined using refractive index measurements.^66, 67^ Urea was titrated into RBD solutions at intervals of 0.2 M, samples were equilibrated for 10 min in between titrations, and the change in spectral data was analyzed using the equation^68^

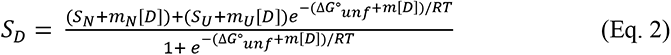

where S_D_ is the signal at a denaturant concentration [D], S_N_ and S_U_ are the signals corresponding to the native and unfolded proteins without denaturant, m_N_ and m_U_ are the slopes of linear dependence of S_N_ and S_U_ with [D], ΔG°_unf_ is the Gibbs free energy change of unfolding, and m is the slope of linear dependence of ΔG_unf_ with [D], R is the universal gas constant, and T is the absolute temperature in Kelvin, respectively.

### ITC binding analysis

ITC was performed on the Malvern Microcal PEAQ-ITC. All experiments were performed in a buffer solution of 50 mM sodium phosphate, 20 mM NaCl, pH 7.0 at 20°C and consisted of eighteen 2 µL injections spaced every 150 seconds. For RBD-ACE2 interactions, ACE2 was set at a concentration of 15 µM while RBD and all variants were injected at 150 µM. For RBD-CC12.1 ScFv interactions, RBD was set at a concentration of 25 µM while CC12.1 was injected at 250 µM. RBD-LY-CoV016 interactions were studied at a concentration of 30 µM RBD and LY-CoV016 was injected at 250 µM. RBD-LY-CoV555, RBD-P2B-2F6, and RBD-REGN10987 ScFv interactions were studied using a RBD concentration of 30 µM, while LY-CoV555, P2B-2F6, and REGN10987 ScFvs were injected at a concentration of 300 µM. All data was collected and analyzed on the Microcal PEAQ-ITC Data Analysis Software. Errors on ΔG and -TΔS were calculated using error propagation formulae.^69^

